# Integrating migratory marine connectivity into shark conservation

**DOI:** 10.1101/2025.10.27.684680

**Authors:** Diego Feitosa Bezerra, Lily Bentley, Ross Dwyer, Dina Nisthar, Anthony J Richardson, Colin Simpfendorfer, Michelle Heupel, Christoph Rohner, Simon Pierce, Daniel Dunn

## Abstract

Understanding migratory connectivity is important for the conservation of highly mobile marine species that face escalating threats across the globe. Establishing baseline information on migratory connectivity is therefore needed to identify regions of conservation focus. Despite efforts to track migratory sharks and rays, information on transboundary movements is limited and often inaccessible to managers and policymakers. Here, we synthesised multimethod movement data for migratory Australian shark and ray species, investigating which species require international engagement to support their population recovery. Based on data from a systematic literature review, we built connectivity networks from telemetry and mark-recapture studies that provide a first baseline for transboundary migratory connectivity for Australian sharks and rays. Of the 31 shark and ray species reviewed, we identified 6 species that link the Australian Exclusive Economic Zone to other national jurisdictions via multispecies migratory connections through the Tasman Sea to New Zealand, through the Tasman and Coral Sea to New Caledonia, and north across the Timor Sea and Torres Strait to Indonesia and Papua New Guinea. White sharks (*Carcharhinus carcharias*) and whale sharks (*Rhincodon typus*) were the most data rich, whereas 14 shark and ray species had no movement information. There is a grave deficiency in available information for endangered or critically endangered migratory shark and ray populations, with 76% having only one or no published studies. This work supports future conservation strategies for migratory sharks that require robust international collaboration and the adoption of integrated and dynamic management approaches.

## Introduction

Migration, the broad-scale movement of populations (Wiens 1997), can transcend ecosystem, national, economic, and cultural boundaries (Harrison et al. 2018; McLean et al. 2023). By connecting distant habitats, migrations contribute to ecosystem function and resilience (Alerstam & Bäckman 2018; Davidson et al. 2020). While migrations can facilitate access to seasonally abundant resources required for key biological processes (Alerstam et al. 2003), they also expose animals to a variety of threats and uncoordinated governance regimes (Dunn et al. 2019; Roberson et al. 2021). This often results in population declines (Teitelbaum & Mueller 2019; Kubelka et al. 2022). Nearly half of all migratory species are in decline, and 20% are at risk of extinction (UNEP-WCMC 2024). Improved management of migratory species is thus urgently needed.

Migratory sharks and rays often exhibit breeding-site philopatry, in which they return to specific regions for parturition or mating after long-distance migrations (Dudgeon et al. 2013; Chapman et al. 2015). These movements link biologically important areas that often span national boundaries, either between adjacent countries or between distant regions connected by extensive transnational migrations (Harrison et al. 2018; Thorburn et al. 2024). This phenomenon, of the geographic linking of individuals and populations across their migratory cycles, is known as migratory connectivity (Webster et al. 2002). Understanding migratory connectivity provides insights into the various stressors an animal encounters during its migration, and how these stressors may accumulate and shape population structure, genetic diversity, and survival rates (Heupel et al. 2015; Somveille et al. 2021; Flack et al. 2022). Therefore, enhancing our knowledge of migratory connectivity underpins the development of effective conservation and management practices aimed at mitigating the decline of migratory species. This is particularly true for elasmobranchs (sharks and rays) – many of which are both highly migratory and under unprecedented threat (Pacoureau et al. 2021; Juan-Jordá et al. 2022), with many populations in decline (Roff et al. 2018; Dulvy et al. 2021; Yan et al. 2021; Simpfendorfer et al. 2023). Globally, approximately one-third of sharks and rays are threatened with extinction (Dulvy et al. 2021). Oceanic populations are more threatened compared to species in other habitats and have declined by 71% over recent decades (Pacoureau et al. 2021).

The Indo-West Pacific, and Australia in particular, is a global biodiversity hotspot for sharks and rays (Pimiento et al. 2023). Australia is frequently cited as a high-priority country for conservation due to its diversity, species richness and the threatened status of its sharks (Dulvy et al. 2017; Stein et al. 2018; Neubauer et al. 2024). Many of these species are dependent on specific habitats in disparate locations and often migrate both within the Australian Exclusive Economic Zone (EEZ) (Stevens et al. 2010; Heupel et al. 2015) and beyond (Chin et al. 2017; McMillan et al. 2019).

Within Australian waters, the Environment Protection and Biodiversity Conservation (EPBC) Act of 1999 is the primary environmental legislation designed to protect and manage flora, fauna, and ecological communities. International cooperation on the conservation of migratory species is coordinated through bilateral and multilateral agreements, and more broadly via the Convention on the Conservation of Migratory Species of Wild Animals (CMS). CMS Appendix I requires binding protection measures with limited scope for exceptions, while Appendix II, where most sharks and rays are listed, only requires cooperation from signatories on management (Lawson & Fordham 2018). The EPBC Act lists eight shark and ray species as threatened, including the whale shark, white shark, grey nurse shark, two river shark species, three sawfish species and Maugean skate (*Zearaja maugean*). The Act further lists 17 species of sharks and rays from the CMS Appendices as migratory, but does not include 12 other CMS-listed species because Australia took a formal exception to those species being listed under CMS (and thus is not compelled to address them as it does other CMS-listed species). While the EPBC Act provides the overarching environmental protection for threatened sharks and rays in Australia, managing their major threat (fishing), including sustainable harvesting and bycatch regulation, is undertaken by state agencies and the Australian Fisheries Management Authority at a national level, and through regional fisheries management organisations (RFMOs) for species caught in shared-stock fisheries in areas beyond national jurisdictions (ABNJ).

Listings of species under CMS and the EPBC Act, and management of sharks and rays by relevant fisheries organisations, are dependent on information on the movement of animals and status of populations being monitored and provided to management and policy fora in an accessible and actionable manner. However, understanding elasmobranch movement at large scales is hampered by major sampling obstacles. Conventional mark-recapture and photo-identification studies are cost-effective and often have large samples of tagged/sighted animals, but the likelihood of recapturing tagged individuals or resighting identifiable individuals is low, and often biased towards highly populated areas and areas of high tourism (Kohler & Turner 2001; Rowat et al. 2009; Mas et al. 2022). Acoustic telemetry can capture local movements, particularly in estuarine or reef ecosystems, but detection probabilities are low across vast areas (Bradford et al. 2019; Spaet et al. 2020). When migratory locations or corridors are unknown, or when species are highly pelagic and travel far from the coastline where receivers are placed, it is usually necessary to undertake expensive satellite tracking or low-resolution archival pop-up light-level tags to capture these movements (Jaine et al. 2014). Despite transnational collaborations expanding acoustic receiver networks, mark-recapture, satellite, and archival pop-up tags remain the main sources of information on large-scale connectivity for elasmobranchs. Thus, difficulties in collecting data on elasmobranch movements persist and have limited our understanding of migration pathways and areas of importance for various life-history stages, which impedes their management and conservation both nationally and internationally. These obstacles to data collection make the available data all the more valuable and increases the importance of aggregating and making use of all available information on shark and ray connectivity.

Here we review and quantify published information on migratory connectivity for shark and ray species listed under the CMS to aggregate evidence of migratory behaviour and transboundary movement requiring international collaboration, and to support more effective management. We build on the Action Plan for Australian Sharks and Rays 2021 that reviewed the extinction risk of all 328 Australian species (Kyne et al. 2021). We conducted a data synthesis of over 30 years of literature on biotelemetry, mark-recapture, and photo-identification studies to establish the first baseline for migratory connectivity of globally at-risk sharks and rays in Australian waters. We also produced network models for migratory shark and ray species, compiling movement information to reveal species-level migratory pathways (Bentley et al. 2024). Our aim is to establish an initial baseline of migratory connectivity for Australian sharks and rays to guide their effective conservation and management. We hope that this will support Australia’s commitments and reporting obligations under international conservation agreements, including the CMS, and the Convention on Biological Diversity. This baseline will determine areas supporting migrations of multiple species, highlight regional and taxonomic data gaps, and help identify nations whose cooperation is needed to improve the conservation and management of individual species.

## Methods

### Species selection criteria

Species selection was based on sharks and rays listed under CMS Appendices I and II, and the Memorandum of Understanding on the Conservation of Migratory Sharks (Shark MoU) (Table 1). The spiny dogfish (*Squalus acanthias*) was not included in our study because only the northern hemisphere populations are listed under CMS. Listing of species under CMS and the Shark MoU is not a systematic process and there are likely to be many species that have not been included yet due to lack of capacity by a CMS member state to nominate them, or lack of available data. We included two further species due to their known large-scale movements and the many individuals tagged in Australasia; the tiger shark (*Galeocerdo cuvier*) and the bull shark (*Carcharhinus leucas*), leading to a potential total of 31 species included in this review.

**Table 1.**
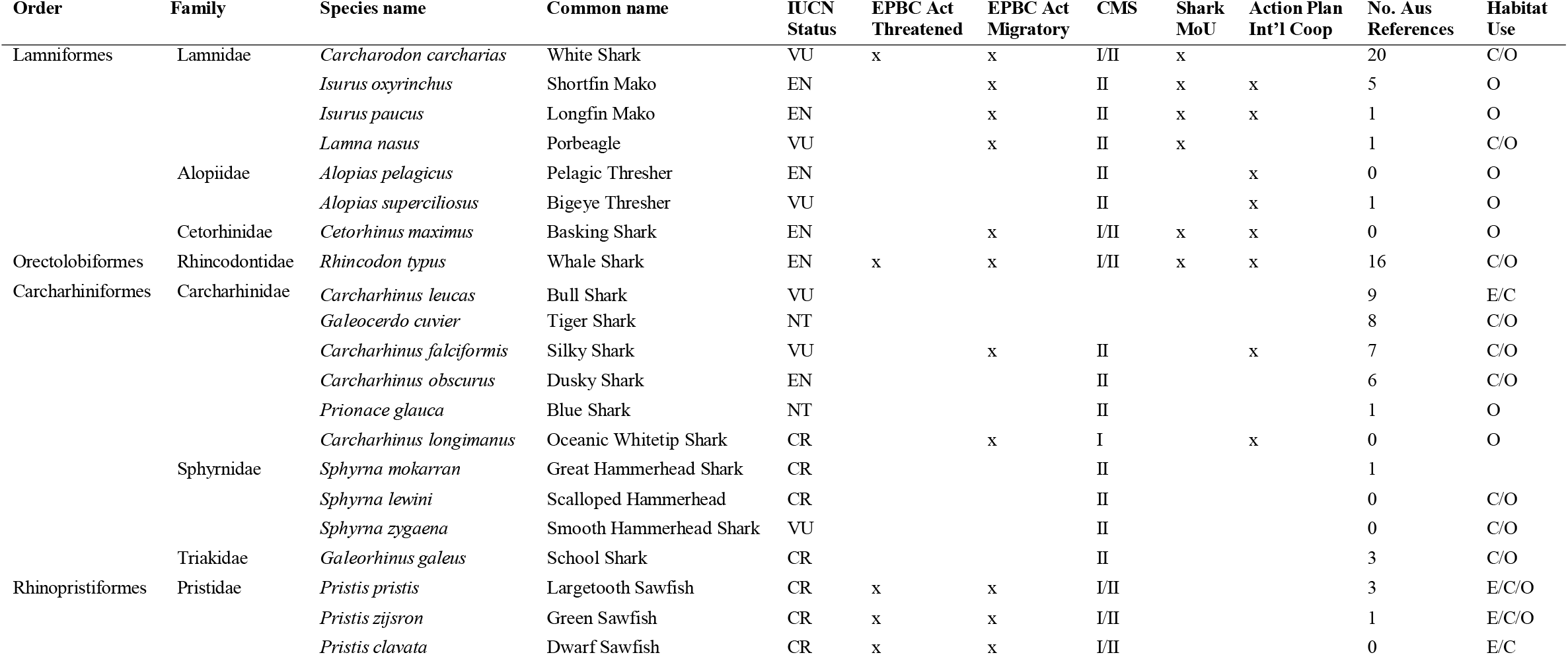
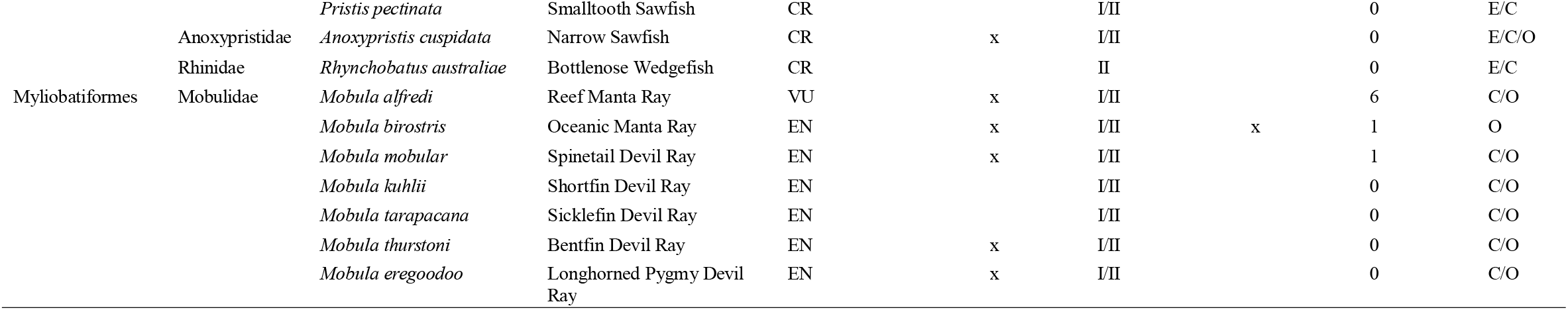
Summary of the 31 species reviewed: their IUCN Red List Categories (Near Threatened (NT), Vulnerable (VU), Endangered (EN), and Critically Endangered (CR)); whether the species is included (marked by an ‘x’) in the Environment Protection and Biodiversity Conservation Act 1999 (EPBC Act), inclusion in the Memorandum of Understanding on the Conservation of Migratory Sharks (Shark MoU) or mentioned as requiring international cooperation in the Action Plan for Australian Sharks and Rays 2021; if and which CMS instrument the species was included in; the total number of references describing connections to Australian water; and the Habitat Use of a species (Euryhaline (E), Coastal (C), or Oceanic (O)).

### Systematic Literature Review Protocol

We followed the PRISMA (Preferred Reporting Items for Systematic Reviews and Meta-Analyses) framework to select and screen peer-reviewed literature for this study (Moher et al. 2009). The PRISMA framework outlines a series of steps, beginning with the identification of relevant studies through comprehensive database searches. Subsequently, references meeting inclusion criteria undergo screening and eligibility assessment based on predefined criteria (here, studies with a migratory connection with Australia). Data are extracted only from selected references (Moher et al. 2009). For this study, we compiled peer-reviewed studies that used biotelemetry, mark-recapture, and photo-identification methods or derived products (e.g., kernel density estimates or state space models) to describe the movements of the 31 candidate species.

All searches for the collection of references were conducted between June and November 2023. We tested the relevance of search strings for a subset of shark and ray species using the Web of Science and Scopus abstract and citation databases (Appendix 1). We noted whether relevant reference (“control”) literature was captured, the total number of references, and how many of the initial papers identified would meet the screening criteria. Within the Web of Science database, all search strings were used to conduct topic searches in English for indexed literature within subscribed databases (WOS, BCI, CCC, DRCI, DIIDW, KJD, MEDLINE, RSCI, SCIELO, ZOOREC), published until June 2023. Within Scopus, all search strings were used to search by keyword, title, and subject area.

### Site aggregations and network models

To determine the connectivity of migratory sharks and rays, the information from the systematic literature review was aggregated into network models following the methods used to develop the MiCO system (www.mico.eco/system; Bentley et al. 2024, Dunn et al. 2019). The general approach is to use network models to aggregate multiple data types describing connectivity. In this approach, individuals reviewing the literature identified areas used by a tagged/visually identified animal (hereafter “sites”), along with “connections” to other sites. The size, period, behaviour, number of animals, sex, and age-class were recorded for each site and connection, if available. The methodology is meant to support description of regional connectivity and sites are categorised into three coarse size classes having 1°, 5° or 10° radii. If no specific animal behaviour was described in the reference reviewed, it was listed as an “observation”.

To generate network models from the many sites and connections identified across various studies, we aggregated overlapping sites into “metasites” (sites made up of multiple sites) and removed studies that appeared to use data employed in other studies. The aggregation and data deduplication process followed Bentley et al. (2024), and has the same four objectives: 1) Integrate data from multiple sites that capture species activity in overlapping locations; 2) Preserve information on large-scale connections (>500 km) resulting from animal movements; 3) Retain high-resolution data on critical reproductive life history stages; and 4) Ensure a minimum sample size of known individuals both within and between sites.

The process of generating network models for shark species entailed five key stages: data preparation, site selection, metasite centroid definition, data duplication checks, and metasite aggregation. In the data preparation stage, sites for each species are compiled and organised based on behaviour, such as reproduction or feeding, as well as factors such as size and number of nearby sites. The ordering of sites ensures that the resolution of reproductive sites is maintained, but that overlapping foraging areas are merged to simplify networks. Subsequently, during site selection, sites are iteratively chosen (i.e., the script examines one at a time based on the ordering) to identify overlapping sites. The metasite centroid definition stage iteratively calculates a new centroid for sites to be aggregated, and then removes any sites whose centroid is no longer within the radii of the new metasite. The centroid is repeatedly recalculated whenever a site is dropped until no further sites are dropped from the metasite. To maintain long distance connections, any sites that have a connection of >500 km were not aggregated. The data duplication stage ensures the integrity of the dataset by checking for similarities such as identical coordinates, route connectivity, authorship, and temporal scope. Duplicate sites informed the location of a centroid of the new metasite but were removed from calculations of the minimum numbers of individuals in a given metasite in the final metasite aggregation stage. Once all sites had been aggregated to a metasite, connections between sites were assigned to their respective metasites, resulting in metaconnections (a connection made up of connections). All code used to develop the network models is available on (see Cover Page below).

We exported the metasites and metaconnections from the MiCO system (www.mico.eco/system), where they are freely available to the public. All subsequent analyses were undertaken in R version 4.3.0, using the packages tidyverse, circlize, ggOceanMaps, jsonlite, and ggExtra (Wickham et al. 2019; Attali & Baker 2022; RTeam et al. 2023). The geodatabase files from the system were used to develop maps of migratory connectivity for all shark and ray species where sufficient information was available. Metasites were also allocated to an EEZ based on the latitude and longitude of the metasite centroid. Adjacency matrices were then constructed from the data on metaconnections that crossed EEZs and were used to generate chord diagrams describing regional connectivity.

## Results

Our data synthesis of 31 shark and ray species that use Australian waters identified 91 relevant peer-reviewed references for 18 species (Table 1, S1). Across these species, we identified 162 metasites and 193 metaconnections based on individual movement data (Table 2). Satellite and acoustic telemetry methods were the primary sources of detailed movement data, with photo-identification and mark-recapture methods providing a minority of evidence. The number of sites and routes identified rose proportionally to tagging efforts and were heavily biased towards iconic species, including white sharks, whale sharks, shortfin mako, tiger sharks and reef manta rays, which accounted for approximately 70% of the sampled individuals.

**Table 2.** The proportion of individuals sampled by sex and life stage, and the number of metasites generated, for the six species demonstrating transboundary movement. The total number of individuals includes those sampled using acoustic and satellite telemetry, and mark-recapture studies.

| Common name | Total number of individuals sampled | Male | Female | Unknown sex | Adult | Subadults | Juvenile | Unknown life stage | Total number of metasites |
| --- | --- | --- | --- | --- | --- | --- | --- | --- | --- |
| White Shark | 917 | 30.8% | 29.1% | 13.5% | 5.4% | 7.7% | 27.2% | 32.9% | 29 |
| Shortfin Mako | 142 | 17.6% | 58.4% | 44.6% | 2.8% | 0 | 96.4% | 1.4% | 16 |
| Whale Shark | 1711 | 36.9% | 11.4% | 51.4% | 0 | 0 | 4.6% | 95.2% | 27 |
| Tiger Shark | 68 | 26.0% | 73.9% | 0 | 30.4% | 2.2% | 64.2% | 3.0% | 28 |
| Blue Shark | 24 | 33.3% | 41.6% | 25% | 29.1% | 0 | 8.3% | 62.5% | 18 |
| School Shark | 57 | 8.7% | 87.7% | 0 | 63.1% | 0 | 0 | 36.8% | 8 |

Thirteen shark and ray species in our review had no published movement information, and another eight had only one relevant publication, which provided movement data specifically within Australian waters (Table 1). Among the 22 endangered species in this study (i.e., those classified as Endangered or Critically Endangered globally by the IUCN Red List; Table 1), 55% (n = 12) had no published movement information. Another five endangered species (23%) had only one published movement study, primarily focused on fine-scale movements in estuaries. Despite the number of data deficient species, our study identified 172 individual sharks of 6 species that engaged in migratory movements either into or out of Australian waters to seven other national jurisdictions and ABNJ (Figure 2).

**Figure 1.**
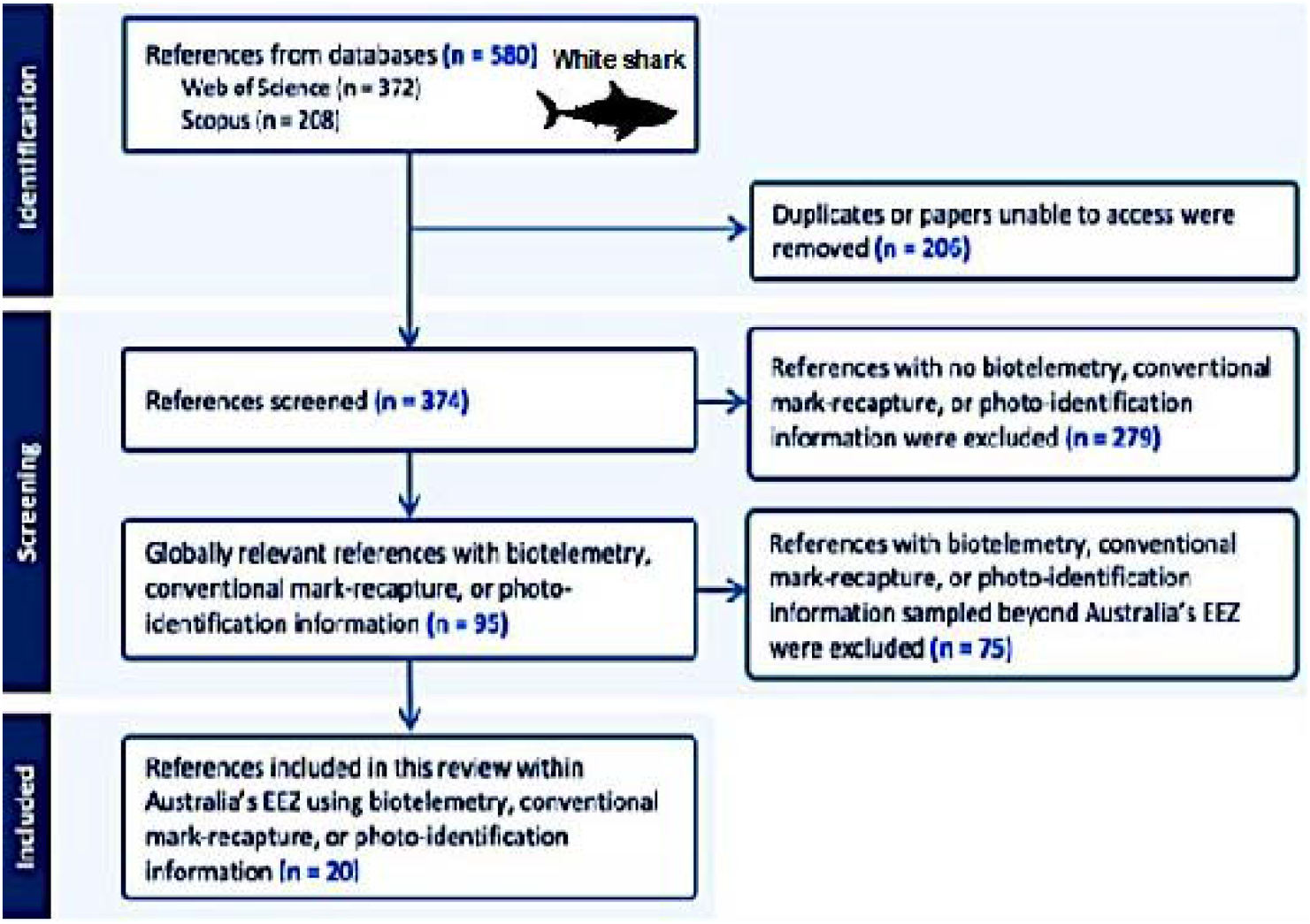
Schematic of the systematic literature review screening protocol illustrated using the white shark (Carcharodon carcharias).

**Figure 2.**
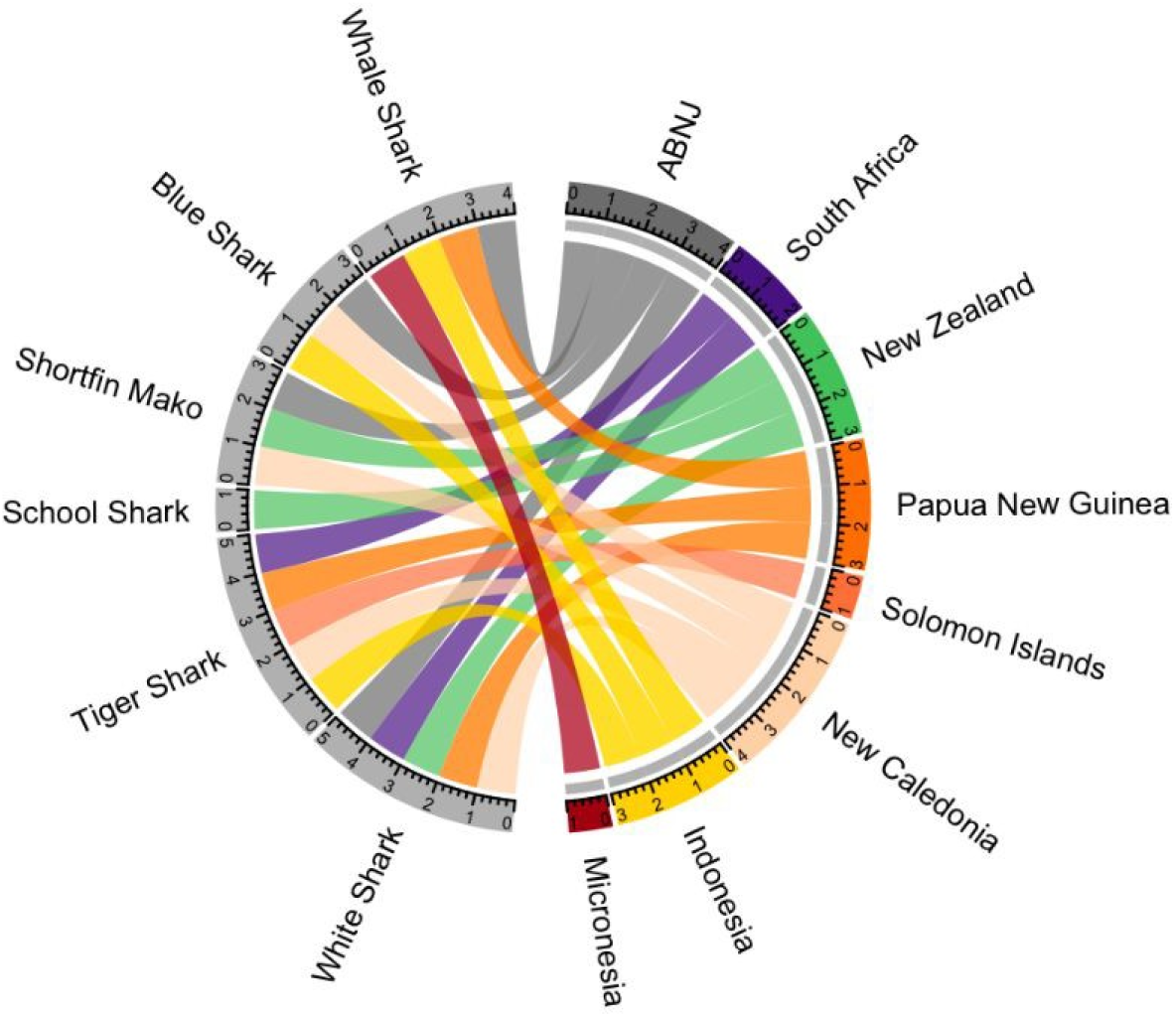
Species with known transboundary movements between the Australian EEZ and other jurisdictions (including areas beyond national jurisdiction; ABNJ). Diagram generated using R package circlize (Gu et al. 2014).

### International transboundary movement of Australian sharks

While we did not find any instances in the published literature of any ray species crossing an international border with Australia, the six migratory shark species in Australian waters that exhibited international transboundary movements were shortfin mako (*Isurus oxyrinchus*), white sharks (*Carcharodon carcharias*), tiger sharks (*Galeocerdo cuvier*), whale sharks (*Rhincodon typus*), blue sharks (*Prionace glauca*) and school sharks (*Galeorhinus galeus*). These species demonstrated complex patterns of habitat use, moving across various habitats, from coastal areas to oceanic habitats, and moving through numerous governance regimes. New Caledonia had the highest number of species with connections shared with Australia, which included the shortfin mako, blue shark, white shark, and tiger shark. White sharks and tiger sharks were the most connected species, with movements linking multiple jurisdictions: New Zealand, New Caledonia, Papua New Guinea, Indonesia, South Africa and Solomon Islands. A total of 34 individual sharks of various species moved between Australia and New Zealand, indicating strong trans-Tasman migratory connectivity.

#### White sharks

For the white sharks, strong white shark connectivity was found between eastern Australia and New Zealand, with 18 individuals migrating to New Zealand and 14 individuals migrating to Australia. White sharks also showed transboundary connectivity between Australia and Papua New Guinea, New Caledonia, and South Africa (Figure 2). Our network models for white shark migratory connectivity show broad-scale movements along the Australian east and west coasts (Figure 3a), and, in general, wide-ranging longitudinal and latitudinal movements.

**Figure 3.**
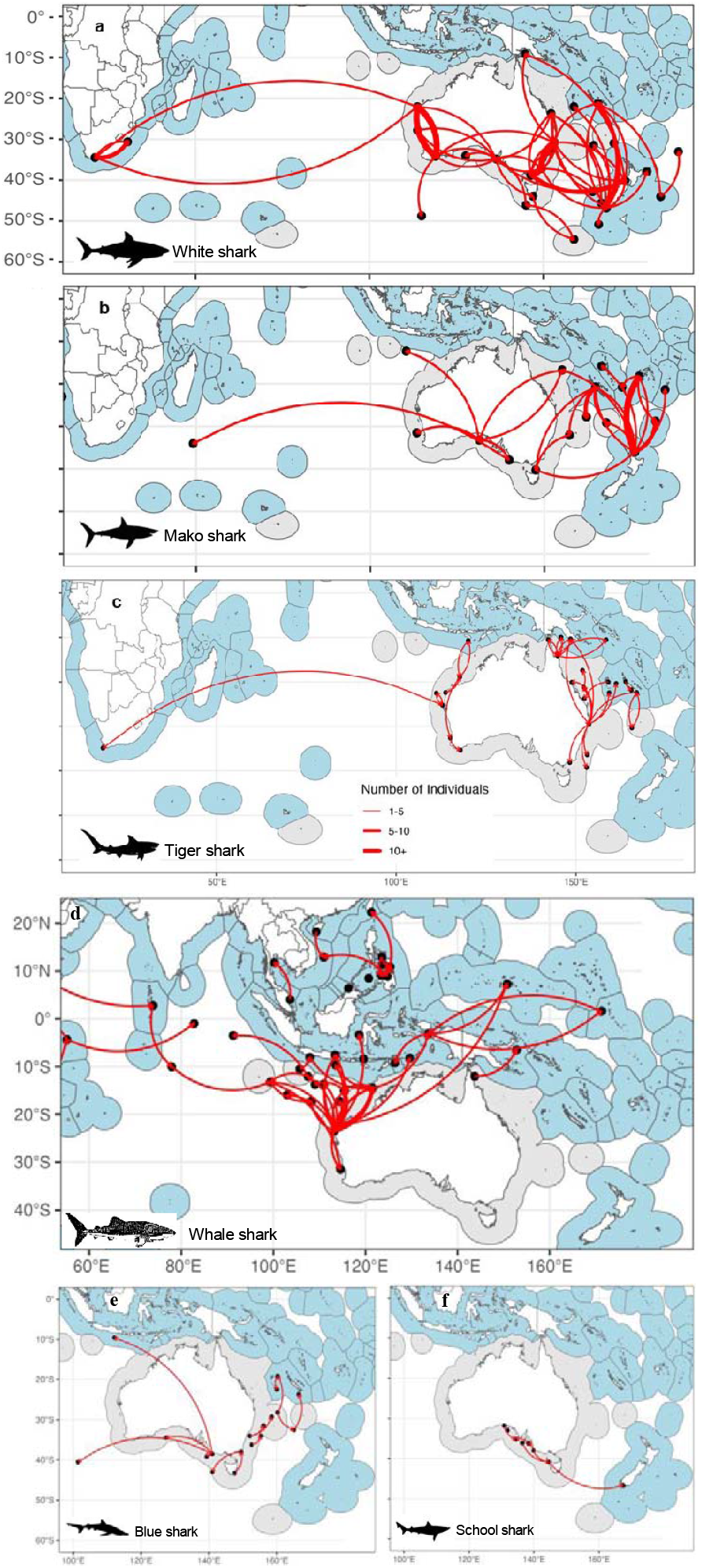
Network connectivity models showing the transboundary movements of six shark species: (a) White shark (b) Shortfin mako, (c) Tiger shark, (d) Whale shark), (e) Blue shark, and (f) School shark. These models depict connectivity across Australia’s Exclusive Economic Zone (EEZ) and into adjacent international waters. Maps were generated using additional data from rnaturalearth (Massicotte and Young, 2023) and EEZ boundaries from Flanders Marine Institute (2023).

#### Shortfin mako

This species demonstrated extensive and diverse movements: some migrated to New Zealand and New Caledonia, while one individual travelled from the Bonney Upwelling Region in southern Australia to the Southwest Indian Ridge in ABNJ. Extensive movements within Australian waters were observed.

#### Tiger sharks

This species frequently traversed coral reef and seagrass ecosystems in coastal environments and demonstrated wide-ranging migratory movements, extending into the Coral Sea region and ABNJ, and crossing multiple national jurisdictions including Papua New Guinea and New Caledonia (Figure 3c).

#### Whale sharks

Published studies on whale shark movements centred around the coastal hotspot of Ningaloo National Marine Park in Western Australia, where they aggregate seasonally to feed (Figure 3d). There were migratory movements into ABNJ of the Indian Ocean, with 27 individuals that exhibited transboundary movements between Australia and Indonesia, Papua New Guinea and overseas Australian territories. Additionally, individuals were recorded migrating from Ningaloo Reef and the Great Barrier Reef to Micronesia.

#### Blue sharks and school sharks

Blue sharks were observed migrating to Indonesia, New Caledonia and ABNJ (Figure 3e). School sharks made extensive long-distance partial migrations using protected pupping areas along the south coast of Australia, with documented movements reaching New Zealand recorded by mark-recapture, acoustic and satellite telemetry methods (Figure 3f).

### Large-scale movements within Australian waters

Movements of acoustically tagged bull sharks (n=73) were restricted to euryhaline and coastal ecosystems, extending from far North Cape York Peninsula to Tollgate Islands in Southeast Australia, and were not recorded moving into other national jurisdictions or ABNJ. Reef manta rays along the Great Barrier Reef in Eastern Australia to northern New South Wales and Ningaloo Marine Park in Western Australia are also well-studied and exhibited well-defined migration patterns that remain within Australian national waters. These populations engaged in seasonal movements between northern and southern reef sections (Figure S1b). Despite these insights, tracking data for other Mobulidae remains limited, with no data on their broader oceanic connectivity and no examples of transboundary movements beyond Australian waters in peer-reviewed literature. A total 473 (out of n = 2478) dusky sharks migrated between western and southern Australian waters, and eight individuals displayed shorter-range movements between Queensland and New South Wales. Whale sharks cross ABNJ to other Australian remote oceanic islands, such as Christmas Island and Cocos Keeling Island, distant from the mainland EEZ.

### Multi-species connections

We identified several key transboundary connections that were important for multiple species. Firstly, the Trans-Tasman pathway represents a key migratory connection for several sharks (white shark, shortfin mako, and school shark). These species migrate across the Great Australian Bight and Bass Strait, extending their migratory pathways through the Tasman Sea to New Zealand. Secondly, shortfin mako, white sharks, and blue sharks shared similar connections extending from the trans-Tasman region to New Caledonia (Figure 3). In addition to these species, tiger sharks exhibited longitudinal migratory pathways further north, transiting the Coral Sea to New Caledonia. Lastly, an area of multispecies migratory connections was found in northern Australia, particularly between Western Australia and Indonesia across the Timor Sea. Both tiger sharks and whale sharks migrated throughout the Timor Sea, travelling from Ningaloo Reef to Indonesia. Moreover, there were connections from waters south of Australia to Indonesia for both shortfin mako and blue sharks.

## Discussion

We provide a baseline for the movement of globally at-risk migratory shark species to support conservation policy in the Indo-West Pacific. Among the nine species requiring international engagement to manage their populations mentioned in The Action Plan for Australian Sharks and Rays (Kyne et al. 2021), only whale sharks and shortfin mako have sufficient data to develop migratory connectivity networks. By synthesising information from 91 published references with movement data from ∼2,900 animals, we identified transboundary movements for six species: white sharks, whale sharks, shortfin mako, blue sharks, school sharks, and tiger sharks. These species visited the EEZs of seven countries or territories, as well as ABNJ. The regional and global scales of these shark migrations necessitate an international collaborative approach to their conservation and management.

### Identification of multispecies migratory connections

The trans-Tasman migratory pathway was identified as important for multiple species in this review. Our network models align with our knowledge of genetic structuring in both white and tiger shark populations across the Tasman Sea and broader West Pacific (Holmes et al. 2017; Bradford et al. 2019). However, genetic information cannot confirm contemporary connectivity or inform spatial management to reduce mortality during migrations, underscoring the importance of collecting individual-based animal movement data and synthesising connectivity data to highlight shared migratory pathways (Figure 3) (Devloo-Delva et al. 2019; McMillan et al. 2019). Multiple species also continued from the Tasman Sea through to New Caledonia. The region has been recognised under the Important Shark and Ray Areas (ISRAs) Subcriterion C4 as a critical movement area linking Norfolk Island and New Caledonia (Jabado et al. 2024). The number of shark species and individuals migrating across the Tasman Sea suggests the need for strengthened bilateral management between Australia and New Zealand.

Multispecies use of the waters from northern Australian through the Timor Sea was also evident. Despite the presence of the Timor Trench, which is up to 3300 m deep and limits the dispersal and connectivity of shark populations (Hirschfeld et al. 2021), both tiger sharks and whale sharks migrated from Ningaloo Reef to Indonesia. Further research could elucidate the degree that the Timor Trench may act as a barrier for movements of coastal sharks, particularly in comparison to the Torres Strait, which may represent a migratory corridor for species that prefer shallow waters but act as a barrier for deep-dwelling species. However, data on movement behaviour in the Timor Sea and Torres Strait are scarce. Although genetic data have provided valuable insights into trans-national elasmobranch populations across the Torres Strait (Chin et al. 2017), tracking data remain limited. There is potential for the movement of multiple species across the strait, as indicated by the distribution, geographical proximity between populations, and genetic studies, such as the critically endangered largetooth sawfish (*Pristis pristis*) (Feutry et al. 2015). Prioritising data collection for poorly studied species in this region is imperative to better understand connectivity, given that Indonesia is one of the world’s largest shark-fishing nations (Sherman et al. 2023). Given the narrow and shallow nature of the Torres Strait, we recommend strategic placement of acoustic receivers throughout this region to capture the potential movement of shallow-obligate sharks and rays moving between Australia and Papua New Guinea. While this approach undertaken in consultation with Traditional Owners and other local stakeholders should work in that region, it is less suitable for the deeper and larger Timor Sea nor for the open ocean areas in the West Pacific. For those regions we recommend deploying satellite and archival tag technology on species caught and retained by fisheries managed through the Western & Central Pacific Fisheries Commission (WCPFC).

We found reduced evidence for multispecies connections with Melanesia and islands east of New Caledonia, though this may be due to lower sampling effort in these regions rather than an actual lack of connectivity. Whale sharks were identified to move from southwest of Papua New Guinea to Kirabati, indicating that while no connection directly from Australian waters was identified, there are movements from the region to the Central Pacific (Womersley et al. 2022). The lack of information on blue shark movement may also be concealing larger-scale movement in this region. Although there were few references in the literature for blue shark in Australian waters, there is extensive information on their ontogenetic habitat shifts across large spatial scales in multiple ocean basins (Maxwell et al. 2019; Fujinami et al. 2021; Madigan et al. 2021; Elliott et al. 2022; Poisson et al. 2024). Similarly, there is a lack of understanding of shortfin mako movements in the region, leading the WCPFC to describe the degree of connectivity as “unclear” and to suggest that the population may comprise “geographically relatively distinct stocks or migratory contingents” (Large et al. 2022). Further research, particularly tracking studies, could address limitations of stock assessments and may provide a more comprehensive understanding of these species’ distributions.

### Transboundary management of migratory sharks

Regional baselines are useful tools for understanding the international transboundary movements of migratory species and, consequently, shared responsibility for the management of migratory populations (Harrison et al. 2018). One existing pathway for multinational collaboration on the conservation of migratory sharks is the CMS and the Sharks Memorandum of Understanding (MoU) that sits beneath it. Of the countries connected by shark movements identified in our study, Australia, New Zealand, New Caledonia (France), and South Africa are signatories to the CMS and the Shark MoU, while Indonesia, Papua New Guinea, the Federated States of Micronesia, and the Solomon Islands are not, nor are they signatories to the Shark MoU. An important step toward better governance of migratory sharks in the region would be for all countries to develop mechanisms to cooperate (e.g., becoming parties to CMS or signatories to the Shark MOU) or establish bilateral agreements where they do not exist. This could then provide political momentum for greater support of research into regional connectivity for sharks and rays. Although the CMS signatory status alone cannot fully address governance gaps for migratory sharks and rays, it plays a constructive role in driving efforts to implement agreed-upon conservation measures.

Effective transboundary management of migratory sharks requires robust data sharing, collaborative research, and policy alignment among nations. Comprehensive data on shark movements, population dynamics, and habitat use are vital to inform management decisions. However, existing research shows a marked bias, with most data focusing on iconic species in wealthier countries, such as Australia, New Zealand, and New Caledonia (France). In contrast, less-studied species in poorer countries often lack sufficient data and have weaker management frameworks. To support the implementation of CMS agreements, it is crucial to prioritise research on the 22 shark and ray species listed under the CMS, many of which currently have few or no connectivity studies in the Indo-West Pacific. With its substantial research capacity and economic resources in the region, Australia is well-placed to lead efforts in addressing these disparities by generating critical data domestically and supporting research initiatives in neighbouring countries with limited resources and capacity. The Australian government could focus on the seventeen shark and ray species listed under the EPBC Act as migratory. Particular attention could be paid to the three sawfish species listed as Threatened under the Act that had only four connectivity studies among them and to hammerhead species. This compares with the other two sharks listed as threatened (white sharks and whale sharks), for which there were 38 studies. An emphasis towards studies that could clarify movement across the Timor Sea and the Torres Strait would also reduce geographic bias towards wealthier regions.

RFMOs play a central role in monitoring and managing the sustainability of fisheries across vast oceanic areas. They implement a variety of measures to mitigate threats to these species, such as setting catch limits, enacting fishery closures, enforcing gear restrictions to reduce bycatch, and designating “no retention” species to prohibit their landing. All West Pacific countries for which transboundary connections with Australia were identified in this study are members of the WCPFC. The WCPFC has a Conservation and Management Measure for Sharks that specifically prohibits: shark finning; retention of whale sharks, oceanic whitetip sharks or silky sharks; the use of shark lines in longlining; and the setting of purse seines on tuna associated with whale sharks (Neubauer et al. 2024). In addition to the Conservation and Management Measure for Sharks, the WCPFC also has a 2021-2030 Shark Research Plan that provides a timeline for assessment of various shark species and a list of prioritised projects (Brouwer & Hamer 2024). Of interest, the Shark Research Plan includes a scoping study to understand stock structure and natal homing across all shark species. Similarly, to address uncertainty in the stock assessment for silky sharks, there is a call for more satellite tagging studies to be undertaken to “resolve fundamental questions… about the degree of natal homing and limited mixing of the stock” (Neubauer et al.2024). For the same reasons, the stock assessment for shortfin mako recommends “additional tagging should be carried out… especially [in] known nursery grounds off southeast Australia and New Zealand, as well as high seas areas to the north and east of New Zealand, where catch-rates are high.” The baseline connectivity models presented here can enhance the understanding of metapopulation dynamics critical to RFMO management, as well as underpin future risk-assessments by allowing cumulative impacts to be accounted for across the known migratory movements of these species. They also provide a framework for how information from various sampling methods can be synthesised to describe regional connectivity for sharks and rays.

It is important to note that while, in this study, we use the term “transboundary” to refer to movements that cross international borders, animals moving across multiple management jurisdictions at any scale face similar challenges. For example, bull sharks demonstrate large movements across Australian state boundaries. As bull shark management is inconsistent across Australian states and territories, these movements are a microcosm of the broader challenge of effectively managing highly migratory species. Further, the scale of these movements could easily have resulted in transboundary movement in regions with more or smaller EEZs. Indeed, while the published movement data for bull sharks and reef manta rays did not show evidence of international transboundary movements, this may be influenced by the limited use of satellite telemetry for bull sharks and/or the predominantly coastal behaviour in reef manta rays (confirmed through aggregated photo-identification data (Armstrong et al. 2019)).

### Using network models to understand migratory connectivity

While information on migratory connectivity can be gleaned from many sampling methods, findings are rarely integrated (Bentley et al. 2024). Our approach identifies sites and connections across methods and integrates them into a single network of the minimum known connectivity for each species. For example, acoustic telemetry data were previously used to describe school shark movement in national waters throughout southern New South Wales, Victoria and eastern Tasmania (Lédée et al. 2021). Our network connectivity models, incorporating satellite telemetry studies, extend these findings by describing transboundary migrations from South Australia to New Zealand waters, and are corroborated by a vast amount of mark-recapture tag data (Hurst et al. 1999). This supports the recent listing of school sharks (also known as tope) under Appendix II of the CMS. Establishing a benchmark for migratory connectivity facilitates international collaboration and supports the identification of interested parties, from national stakeholders to intergovernmental organisations, with management mandates related to sharks and rays. These benchmarks also identify the regions and species for which data gaps persist, providing evidence to guide the prioritisation of future research efforts. Benchmarks for well-studied species (e.g., the white shark in southeastern Australia) also demonstrate the power and feasibility of this method. The approach adopted here offers a transferable model to help develop baselines for migratory connectivity in sharks and rays in other regions, which is critical to conserving these threatened taxa.

Some caveats should also be considered when interpreting our results. Network models demonstrate movement patterns from known data, but also reflect biases in the location of tagging or acoustic receivers, which can potentially skew results. Our results mostly relied on satellite telemetry data and passive acoustic telemetry studies, methods that are commonly deployed in coastal regions (Hussey et al. 2015). Another key limitation is that network models do not capture the precise routes individuals take between nodes. Connections between nodes represent observed migratory links, not a precise movement path. As such, the models provide a simplified representation of complex migratory behaviours and cannot readily inform spatial management of migratory corridors without further interrogation of the underlying data.

It is important to acknowledge that unpublished data, including telemetry studies not yet available in the public domain, were not included here but could provide additional insights into migratory patterns and further inform our understanding of species connectivity. Many research efforts are never published, or are published in un-indexed grey literature that was not reviewed in this study, potentially resulting in apparent “gaps” in which studies have indeed been undertaken. As such, this study should be viewed as an assessment of the minimum known transboundary connectivity for Australian sharks and rays, and further work should be undertaken to incorporate any available information on shark and ray movements. Together with improvements in spatial resolution and gap-filling to address geographic biases, these measures could offer a more comprehensive view of shark and ray migratory connectivity.

Further tagging efforts are needed to address the significant data gaps that persist within Australia and neighbouring countries. Critically, given the transboundary nature of many shark and ray migrations and distributions, regional funding to raise capacity across all neighbouring countries is essential to support Australian shark and ray populations and their important habitats. Our synthesis of available evidence identifies several key opportunities for international partnerships to strengthen migratory sharks and ray conservation in the Indo-West Pacific. Of particular interest are multispecies migratory connections between countries shared by multiple shark and ray species, such as Australia and Indonesia, Papua New Guinea, and New Zealand. Aggregation of such shared connections in unified models of migratory connectivity could help foster the cooperative transboundary management required to sustain shark and ray populations.

## References

Alerstam T, Bäckman J. 2018. Ecology of animal migration. Current Biology.

Alerstam T, Hedenström A, Åkesson S. 2003. Long-distance migration: Evolution and determinants. Oikos.

Armstrong AO, Armstrong AJ, Bennett MB, Richardson AJ, Townsend KA, Dudgeon CL. 2019. Photographic identification and citizen science combine to reveal long distance movements of individual reef manta rays Mobula alfredi along Australia’s east coast. Marine Biodiversity Records 12.

Attali D, Baker C. 2022. ggExtra: Add Marginal Histograms to “ggplot2”, and More “ggplot2” Enhancements. The R Journal.

Bentley L et al. 2024, June 12. Marine megavertebrate migrations connect the global oceans. Available from https://www.researchsquare.com/article/rs-4457815/v1.

Bradford BD, Harasti D, Lee K, Gallen C, Bradford R. 2019. Broad-scale movements of juvenile white sharks Carcharodon carcharias in eastern Australia from acoustic and satellite telemetry. Marine Ecology Progress Series 619.

Brouwer S, Hamer P. 2024. The Commission for the Conservation and Management of Highly Migratory Fish Stocks in the Western and Central Pacific Ocean. Scientific Committee Twentieth Regular Session. WCPFC-SC20-2024:1–251.

Chapman DD, Feldheim KA, Papastamatiou YP, Hueter RE. 2015. There and back again: A review of residency and return migrations in sharks, with implications for population structure and management. Annual Review of Marine Science 7.

Chin A, Simpfendorfer CA, White WT, Johnson GJ, McAuley RB, Heupel MR. 2017. Crossing lines: A multidisciplinary framework for assessing connectivity of hammerhead sharks across jurisdictional boundaries. Scientific Reports 7.

Davidson SC et al. 2020. Ecological insights from three decades of animal movement tracking across a changing Arctic. Science 370.

Dudgeon CL, Lanyon JM, Semmens JM. 2013. Seasonality and site fidelity of the zebra shark, Stegostoma fasciatum, in southeast Queensland, Australia. Animal Behaviour 85:471–481.

Dulvy NK et al. 2021. Overfishing drives over one-third of all sharks and rays toward a global extinction crisis. Current Biology 31.

Dulvy NK, Simpfendorfer CA, Davidson LNK, Fordham S V., Bräutigam A, Sant G, Welch DJ. 2017. Challenges and Priorities in Shark and Ray Conservation.

Dunn DC et al. 2019. The importance of migratory connectivity for global ocean policy. Proceedings of the Royal Society B: Biological Sciences 286.

Elliott RG, Montgomery JC, Penna A Della, Radford CA. 2022. Satellite tags describe movement and diving behaviour of blue sharks Prionace glauca in the southwest Pacific. Marine Ecology Progress Series 689:77–94.

Feutry P, Kyne PM, Pillans RD, Chen X, Marthick JR, Morgan DL, Grewe PM. 2015. Whole mitogenome sequencing refines population structure of the Critically Endangered sawfish Pristis pristis. Marine Ecology Progress Series 533.

Flack A et al. 2022. New frontiers in bird migration research.

Fowler S. 2014. The Conservation Status of Migratory Sharks. UNEP/CMS Secretariat, Memorandum of Understanding on the Conservation of Migratory Sharks.

Fujinami Y, Shiozaki K, Hiraoka Y, Semba Y, Ohshimo S, Kai M. 2021. Seasonal migrations of pregnant blue sharks Prionace glauca in the northwestern Pacific. Marine Ecology Progress Series 658.

Harrison AL et al. 2018. The political biogeography of migratory marine predators. Nature Ecology and Evolution 2.

Heupel MR, Simpfendorfer CA, Espinoza M, Smoothey AF, Tobin A, Peddemors V. 2015. Conservation challenges of sharks with continental scale migrations. Frontiers in Marine Science 2.

Hirschfeld M, Dudgeon C, Sheaves M, Barnett A. 2021. Barriers in a sea of elasmobranchs: From fishing for populations to testing hypotheses in population genetics.

Holmes BJ, Williams SM, Otway NM, Nielsen EE, Maher SL, Bennett MB, Ovenden JR. 2017. Population structure and connectivity of tiger sharks (Galeocerdo cuvier) across the Indo-Pacific Ocean basin. Royal Society Open Science 4.

Hurst RJ, Baglet NW, McGregor GA, Francis MP. 1999. Movements of the New Zealand school shark, Galeorhinus galeus, from tag returns. New Zealand Journal of Marine and Freshwater Research 33:29–48.

Hussey NE et al. 2015. Aquatic animal telemetry: A panoramic window into the underwater world. Science 348.

Jaine FRA, Rohner CA, Weeks SJ, Couturier LIE, Bennett MB, Townsend KA, Richardson AJ. 2014. Movements and habitat use of reef manta rays off eastern Australia: Offshore excursions, deep diving and eddy affinity revealed by satellite telemetry. Marine Ecology Progress Series 510.

Juan-Jordá MJ, Murua H, Arrizabalaga H, Merino G, Pacoureau N, Dulvy NK. 2022. Seventy years of tunas, billfishes, and sharks as sentinels of global ocean health. Science 378.

Kohler NE, Turner PA. 2001. Shark tagging: A review of conventional methods and studies.

Kubelka V, Sandercock BK, Székely T, Freckleton RP. 2022. Animal migration to northern latitudes: environmental changes and increasing threats.

Kyne PM, Heupel MR, White WT, Simpfendorfer C. 2021. The Action Plan for Australian Sharks and Rays 2021. National Environmental Research Program Marine Biodiversity Hub. 1 ed.

Large K, Neubauer P, Brouwer S. 2022. SCIENTIFIC COMMITTEE EIGHTTEENTH REGULAR SESSION ELECTRONIC MEETING Stock assessment of Southwest Pacific Shortfin Mako shark Dragonfly Data Science 2 Saggitus Consulting Stock assessment of Southwest Pacific Shortfin Mako shark.

Lédée EJI et al. 2021. Continental-scale acoustic telemetry and network analysis reveal new insights into stock structure. Fish and Fisheries 22.

Madigan DJ, Shipley ON, Carlisle AB, Dewar H, Snodgrass OE, Hussey NE. 2021. Isotopic Tracers Suggest Limited Trans-Oceanic Movements and Regional Residency in North Pacific Blue Sharks (Prionace glauca). Frontiers in Marine Science 8.

Mas F, Cortés E, Coelho R, Defeo O, Forselledo R, Jiménez S, Miller P, Domingo A. 2022. Shedding rates and retention performance of conventional dart tags in large pelagic sharks: Insights from a double-tagging experiment on blue shark (Prionace glauca). Fisheries Research 255.

Maxwell SM, Scales KL, Bograd SJ, Briscoe DK, Dewar H, Hazen EL, Lewison RL, Welch H, Crowder LB. 2019. Seasonal spatial segregation in blue sharks (Prionace glauca) by sex and size class in the Northeast Pacific Ocean. Diversity and Distributions 25.

McLean M, Warner B, Markham R, Fischer M, Walker J, Klein C, Hoeberechts M, Dunn DC. 2023. Connecting conservation & culture: The importance of Indigenous Knowledge in conservation decision-making and resource management of migratory marine species. Marine Policy 155.

McMillan MN, Huveneers C, Semmens JM, Gillanders BM. 2019. Partial female migration and cool-water migration pathways in an overfished shark. ICES Journal of Marine Science 76.

Moher D et al. 2009. Preferred reporting items for systematic reviews and metaanalyses: The PRISMA statement.

Neubauer P, Kim K, Large K, Brouwer S. 2024. SCIENTIFIC COMMITTEE TWENTIETH REGULAR SESSION Manila, Philippines Stock Assessment of Silky Shark in the Western and Central Pacific Ocean 2024.

Pacoureau N et al. 2021. Half a century of global decline in oceanic sharks and rays. Nature 589.

Pacoureau N et al. 2023. Conservation successes and challenges for wide-ranging sharks and rays. Proceedings of the National Academy of Sciences of the United States of America 120.

Pimiento C, Albouy C, Silvestro D, Mouton TL, Velez L, Mouillot D, Judah AB, Griffin JN, Leprieur F. 2023. Functional diversity of sharks and rays is highly vulnerable and supported by unique species and locations worldwide. Nature Communications 14.

Poisson F, Demarcq H, Coudray S, Bohn J, Camiñas JA, Groul JM, March D. 2024. Movement pathways and habitat use of blue sharks (Prionace glauca) in the Western Mediterranean Sea: Distribution in relation to environmental factors, reproductive biology, and conservation issues. Fisheries Research 270.

Roberson LA et al. 2021. Multinational coordination required for conservation of over 90% of marine species. Global Change Biology 27.

Roff G, Brown CJ, Priest MA, Mumby PJ. 2018. Decline of coastal apex shark populations over the past half century. Communications Biology 1.

Rowat D, Speed CW, Meekan MG, Gore MA, Bradshaw CJA. 2009. Population abundance and apparent survival of the Vulnerable whale shark Rhincodon typus in the Seychelles aggregation. Page ORYX.

Runge CA, Gallo-Cajiao E, Carey MJ, Garnett ST, Fuller RA, McCormack PC. 2017. Coordinating Domestic Legislation and International Agreements to Conserve Migratory Species: A Case Study from Australia. Conservation Letters 10.

Sherman CS, Digel ED, Zubick P, Eged J, Haque AB, Matsushiba JH, Simpfendorfer CA, Sant G, Dulvy NK. 2023. High overexploitation risk due to management shortfall in highly traded requiem sharks. Conservation Letters 16.

Simpfendorfer CA et al. 2023. Widespread diversity deficits of coral reef sharks and rays. Science 380.

Somveille M, Bay RA, Smith TB, Marra PP, Ruegg KC. 2021. A general theory of avian migratory connectivity. Ecology Letters 24.

Spaet JLY, Patterson TA, Bradford RW, Butcher PA. 2020. Spatiotemporal distribution patterns of immature Australasian white sharks (Carcharodon carcharias). Scientific Reports 10.

Stein RW, Mull CG, Kuhn TS, Aschliman NC, Davidson LNK, Joy JB, Smith GJ, Dulvy NK, Mooers AO. 2018. Global priorities for conserving the evolutionary history of sharks, rays and chimaeras. Nature Ecology and Evolution 2.

Stevens JD, Bradford RW, West GJ. 2010. Satellite tagging of blue sharks (Prionace glauca) and other pelagic sharks off eastern Australia: Depth behaviour, temperature experience and movements. Marine Biology 157.

Team RC. 2023. R Core Team 2023 R: A language and environment for statistical computing. R foundation for statistical computing. https://www.R-project.org/. R Foundation for Statistical Computing.

Teitelbaum CS, Mueller T. 2019. Beyond Migration: Causes and Consequences of Nomadic Animal Movements.

Thorburn J et al. 2024. Assessing the potential of acoustic telemetry to underpin the regional management of basking sharks (Cetorhinus maximus). Animal Biotelemetry 12. BioMed Central Ltd.

UNEP-WCMC. 2024. State of the World’s Migratory Species. Convention on the Conservation of Migratory Species of Wild Animals:1–88. Available from https://www.unep-wcmc.org.

Webster MS, Marra PP, Haig SM, Bensch S, Holmes RT. 2002. Trends in Ecology & Evolution 17.

Wickham H et al. 2019. Welcome to the Tidyverse. Journal of Open Source Software 4.

Wiens JA. 1997. Metapopulation Dynamics and Landscape Ecology. Page Metapopulation Biology.

Womersley FC et al. 2022. Global collision-risk hotspots of marine traffic and the world’s largest fish, the whale shark. Proceedings of the National Academy of Sciences of the United States of America 119.

Yan HF, Kyne PM, Jabado RW, Leeney RH, Davidson LNK, Derrick DH, Finucci B, Freckleton RP, Fordham S V., Dulvy NK. 2021. Overfishing and habitat loss drives range contraction of iconic marine fishes to near extinction. Science Advances 7.

