## Supplementary material for "Integrating migratory marine connectivity into shark conservation": Table S1 S2 (from another version of this preprint: version 1, 2025-10-27)

**Supplemental Information**

**Appendix 1: Screening process and search strings**

**Papers were extracted if:**
Includes spatial information through the relevant methods about the migratory shark and ray species list within Australia EEZ **(Figure 1)**. "Relevant" references were identified as:

1) Relevant if: (a & b & c)
   a) Includes species in the search string, AND
   b) Includes geographic information, AND
   c) The research methods include satellite and/or acoustic telemetry, mark-recapture, or photo-ID.

2) Non-relevant if: (a & b & c & d)
   a) Does not include species that were used in the search string OR
   b) Does not include any geographic information for the study OR
   c) Does not use satellite and/or acoustic telemetry, mark-recapture or photo-ID in the research methods OR Reference was not in English, or a copy could not be obtained.

**Search strings:**
Web of Science
ALL=((<Genus species> OR <Common name>) AND (eDNA OR isotop* OR genetic* OR mark* OR recap* OR argos OR telemetry OR sonic* OR tag* OR gps* OR acoustic* OR pat* OR conventional OR geolocat* OR ( photo AND id ) ) AND ( migrat* OR connect* OR move* OR feed* OR forag* OR breed* OR dispers* OR aggregat* OR site* OR corridor* OR route* OR track* OR winter* OR habitat* OR biogeograph* OR ( population AND structure)))

SCOPUS
TITLE-ABS-KEY (("Genus species" OR "Common name") AND ( edna OR isotop* OR genetic* OR mark* OR sonic* OR recap* OR argos OR telemetry OR tag* OR gps* OR acoustic* OR pat* OR conventional OR geolocat* OR ( photo AND id ) ) AND ( migrat* OR connect* OR move* OR feed* OR forag* OR breed* OR dispers* OR aggregat* OR site* OR corridor* OR route* OR track* OR winter* OR habitat* OR biogeograph* OR ( population AND structure)))

**Table S1.** Summary of the 29 CMS-listed species reviewed: their IUCN Red List Categories (Near Threatened (NT), Vulnerable (VU), Endangered (EN), and Critically Endangered (CR)); whether the species is included (marked by an 'x') in the Environment Protection and Biodiversity Conservation Act 1999 (EPBC Act), inclusion in the Memorandum of Understanding on the Conservation of Migratory Sharks (Shark MoU) or mentioned as requiring international cooperation in the Action Plan for Australian Sharks and Rays 2021; if and which CMS instrument the species was included in; the total number of references describing connections to Australian water; and the Habitat Use of a species (Euryhaline (E), Coastal (C), or Oceanic (O)).

| Order | Family | Species name | Common name | IUCN Status | EPBC Act Threatened | EPBC Act Migratory | CMS | Shark MoU | Action Plan Int'l Coop | No. Aus References | Habitat Use |
|---|---|---|---|---|---|---|---|---|---|---|---|
| Lamniformes | Lamnidae | *Carcharodon carcharias* | White Shark | VU | x | x | I/II | x | | 20 | C/O |
| | | *Isurus oxyrinchus* | Shortfin Mako | EN | | x | II | x | x | 5 | O |
| | | *Isurus paucus* | Longfin Mako | EN | | x | II | x | x | 0 | O |
| | | *Lamna nasus* | Porbeagle | VU | | x | II | x | | 0 | C/O |
| | Alopiidae | *Alopias pelagicus* | Pelagic Thresher | EN | | | II | | x | 0 | O |
| | | *Alopias superciliosus* | Bigeye Thresher | VU | | | II | | x | 1 | O |
| | Cetorhinidae | *Cetorhinus maximus* | Basking Shark | EN | | x | I/II | x | x | 0 | O |
| Orectolobiformes | Rhincodontidae | *Rhincodon typus* | Whale Shark | EN | x | x | I/II | x | x | 16 | C/O |
| Carcharhiniformes | Carcharhinidae | *Carcharhinus falciformis* | Silky Shark | VU | | x | II | | x | 0 | C/O |
| | | *Carcharhinus obscurus* | Dusky Shark | EN | | | II | | | 6 | C/O |
| | | *Prionace glauca* | Blue Shark | NT | | | II | | | 2 | O |
| | | *Carcharhinus longimanus* | Oceanic Whitetip Shark | CR | | x | I | | x | 0 | O |
| | Sphyrnidae | *Sphyrna mokarran* | Great Hammerhead | CR | | | II | | | 1 | C |
| | | *Sphyrna lewini* | Scalloped Hammerhead | CR | | | II | | | 0 | C/O |
| | | *Sphyrna zygaena* | Smooth Hammerhead | VU | | | II | | | 0 | C/O |
| | Triakidae | *Galeorhinus galeus* | School Shark | CR | | | II | | | 3 | C/O |
| Rhinopristiformes | Pristidae | *Pristis pristis* | Largetooth Sawfish | CR | x | x | I/II | | | 3 | E/C/O |
| | | *Pristis zijsron* | Green Sawfish | CR | x | x | I/II | | | 0 | E/C/O |
| | | *Pristis clavata* | Dwarf Sawfish | CR | x | x | I/II | | | 0 | E/C |
| | | *Pristis pectinata* | Smalltooth Sawfish | CR | | | I/II | | | 0 | E/C |
| | Anoxypristidae | *Anoxypristis cuspidata* | Narrow Sawfish | CR | | x | I/II | | | 0 | E/C/O |
| | Rhinidae | *Rhynchobatus australiae* | Bottlenose Wedgefish | CR | | | II | | | 0 | E/C |
| Myliobatiformes | Mobulidae | *Mobula alfredi* | Reef Manta Ray | VU | | x | I/II | | | 6 | C/O |

| | | | | | | | |
|---|---|---|---|---|---|---|---|
| *Mobula birostris* | Oceanic Manta Ray | EN | x | I/II | x | 1 | O |
| *Mobula mobular* | Spinetail Devil Ray | EN | x | I/II | | 0 | C/O |
| *Mobula kuhlii* | Shortfin Devil Ray | EN | | I/II | | 0 | C/O |
| *Mobula tarapacana* | Sicklefin Devil Ray | EN | | I/II | | 0 | C/O |
| *Mobula thurstoni* | Bentfin Devil Ray | EN | x | I/II | | 0 | C/O |
| *Mobula eregoodoo* | Longhorned Pygmy Devil Ray | EN | x | I/II | | 0 | C/O |

**Table S2** Results from the data-rich, unlisted species (tiger sharks and bull sharks) case study including the number of references including information in Australian waters, metasites and metaconnections, and the proportion of individuals sampled by sex and life stage.

| Common name | Aus Refs | Total metasites | Total meta-connections | Female | Unknown sex | Adult | Juvenile | Unknown life stage | Transboundary movement |
|---|---|---|---|---|---|---|---|---|---|
| Tiger Shark | 8 | 27 | 31 | 73.9% | 0 | 30.4% | 66.4% | 3.0% | Yes |
| Bull Shark | 9 | 6 | 7 | 62.92% | 11.05% | 67.41% | 12.63% | 19.94% | No |
