## Supplementary material for "Integrating migratory marine connectivity into shark conservation": Figure S1 S2 S3a S3b (from another version of this preprint: version 1, 2025-10-27)

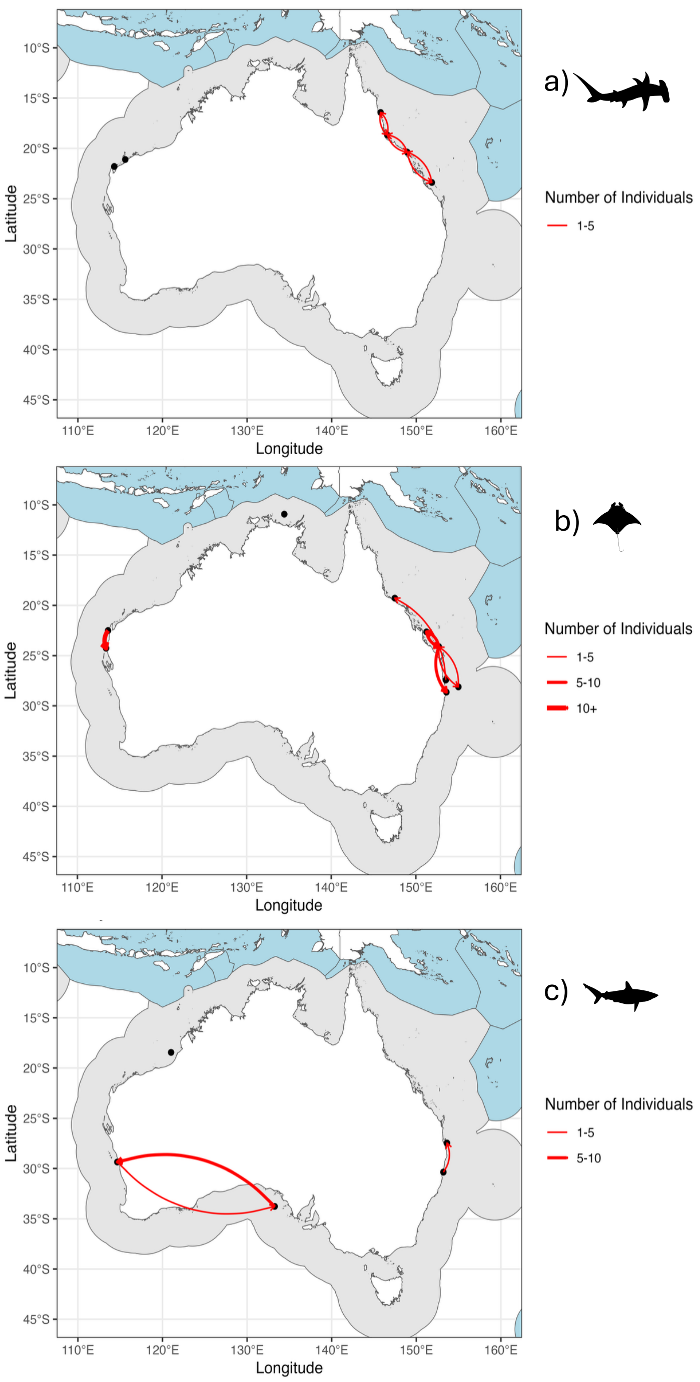


***Figure S.1 -*** *Network connectivity models showing the movements of shark and ray species that did not cross Australian boundaries: (a) Great hammerhead, (b) Reef manta ray, (c) Dusky shark.*

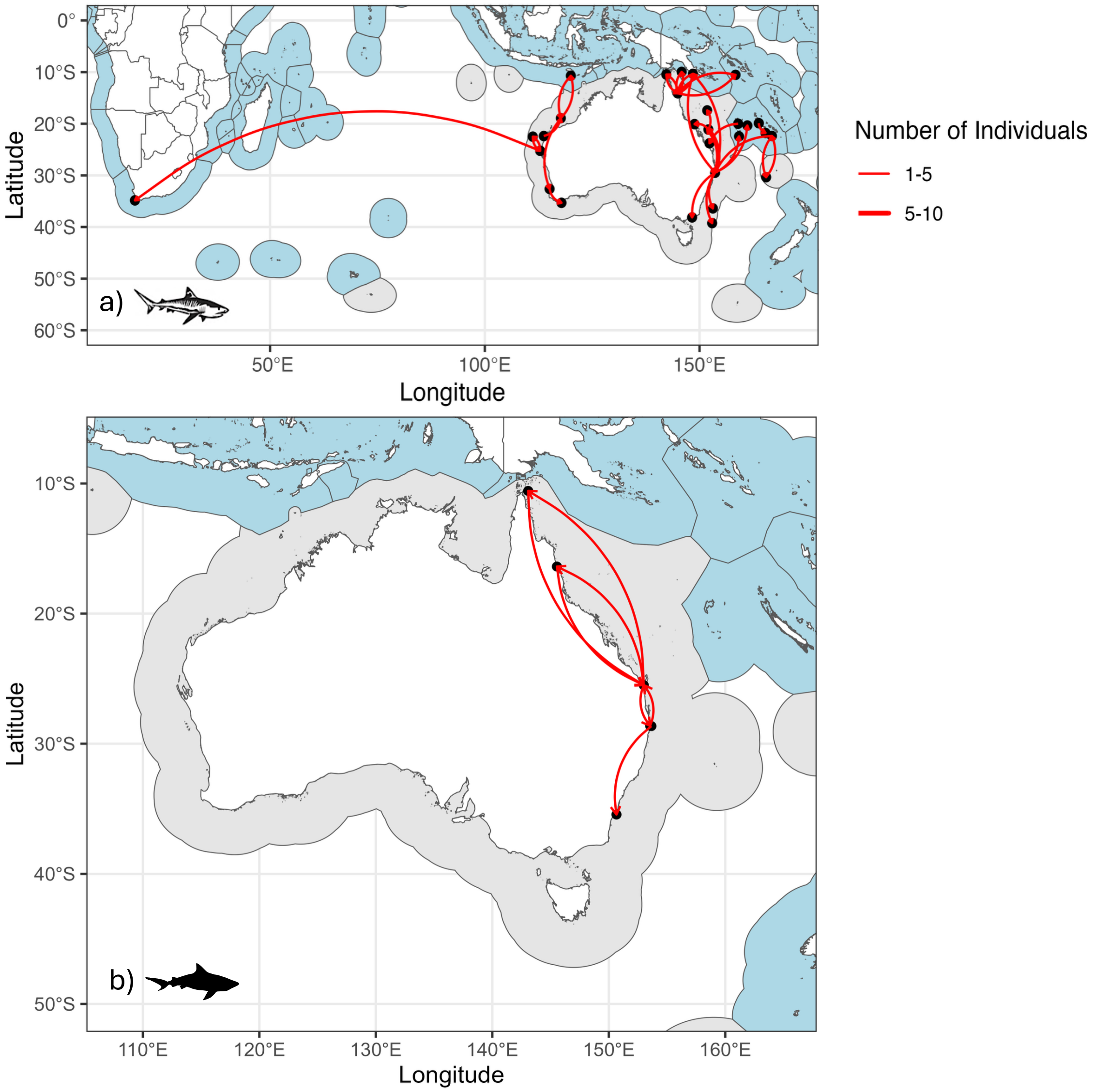


***Figure S.2 -*** *Network connectivity models showing the movements of two well-studied species that are currently not listed under the CMS: (a) tiger shark; (b) bull shark.*

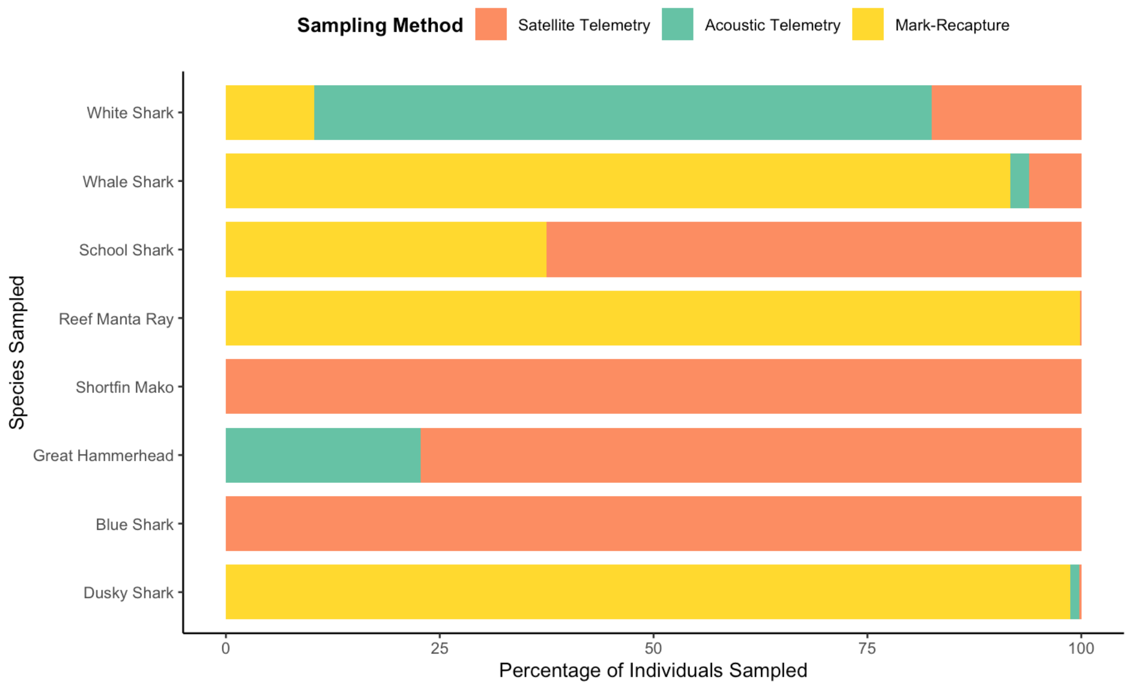


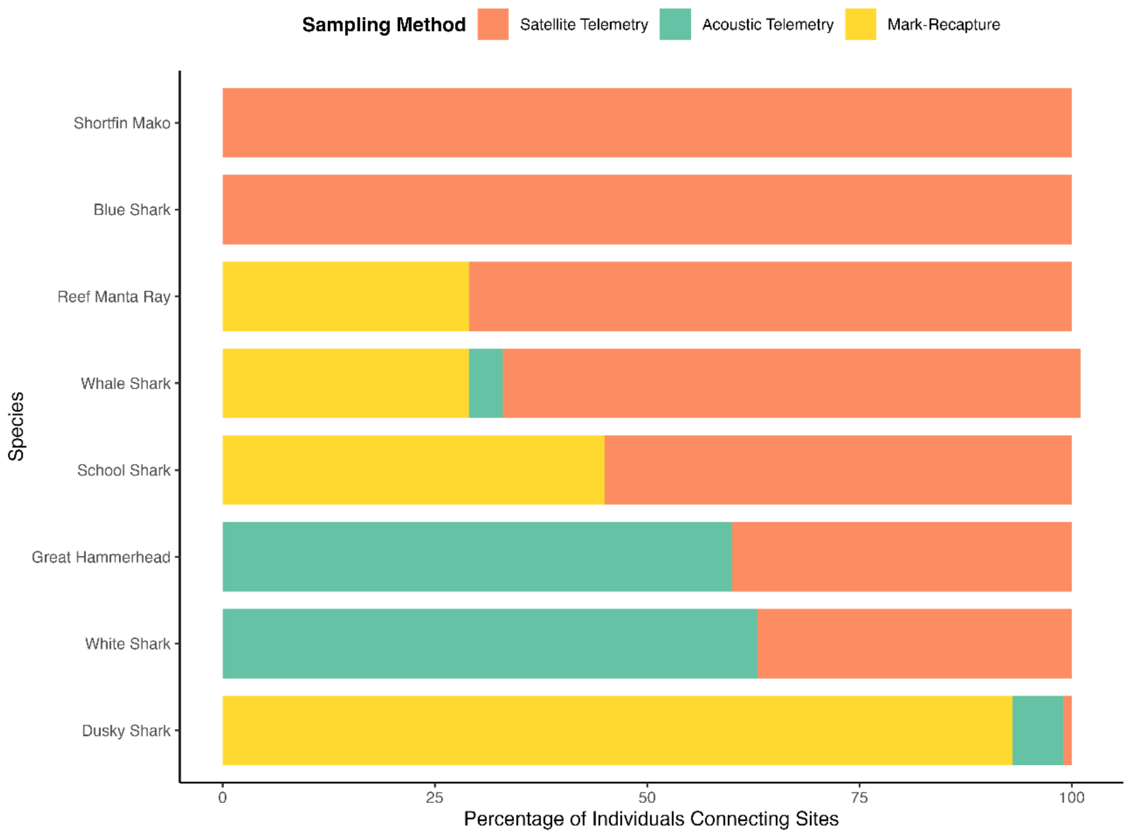


***Figure S.3 a)*** *The percentage of individuals sampled per method and species.* ***b)*** *The percentage of individuals connecting sites per method and species. Mark-Recapture represents conventional tagging and photo-ID combined.*
